# Skull Bone Marrow Drainage and Its Associations with Inflammation, Sleep Quality, and Cognitive Performance

**DOI:** 10.1101/2025.11.10.687593

**Authors:** Ying Zhou, Haidi Jin, Xiao Zhu, Yifei Li, Ziyu Zhou, Xin Huang, Huihong Ke, Mengmeng Fang, Jianzhong Sun, Min Lou

## Abstract

**Background:** Recent studies using human craniotomy samples and murine models have revealed channels that connect the inner skull cortex to the skull marrow cavities. While cerebrospinal fluid (CSF) tracer access to skull bone marrow has been described in CSF circulation disorders, evidence on regional patterns and clinical correlates in broader clinical cohorts is limited.

**Methods:** Three-dimensional imaging was performed at baseline, 4.5, 15, and 39 hours post-gadolinium administration in 87 patients with neurological conditions. Signal changes in skull bone marrow near the superior sagittal sinus, lateral fissure, and cisterna magna were recorded. CSF drainage function was characterized as the percentage changes in the signal unit ratio within the skull bone marrow from baseline to time points post administration of gadolinium.

**Results:** The tracer drained from CSF into the skull bone marrow, highlighting the drainage variations nearby different regions. Reduced drainage was associated with female sex, hypertension, diabetes, and elevated neutrophil levels (*p* < 0.05). Drainage near the superior sagittal sinus was inversely associated with sleep quality and positively associated with cognitive scores, and partially accounted for the association of sleep quality with cognitive function (indirect effect Δβ = −0.092; bootstrap 95% CI -0.195 to -0.004; proportion mediated ≈ 38%).

**Conclusion:** These results support the presence of skull bone marrow adjacent routes for CSF tracer movement and suggest that skull bone marrow tracer dynamics relate to systemic inflammation, sleep quality, and cognitive performance in this clinical cohort.

## Introduction

The traditional belief that the central nervous system (CNS) is immune-privileged due to the protection offered by the intact blood-brain barrier has been challenged by recent research breakthroughs. Emerging evidence indicates that neuroimmune interactions extend beyond the confines of the CNS. Notably, the discovery of lymphatic vessels within the dura mater, which are responsible for directly clearing substances and brain-specific solutes from the cerebrospinal fluid (CSF), significantly alters our previous understanding ^1-3^. Recent studies using human craniotomy samples and murine models have revealed channels that connect the inner skull cortex to the skull marrow cavities ^4^. These channels, known as skull-meninges connections (SMCs), create a direct pathway for immune cells moving to the dura mater from the adjacent skull bone marrow ^5^.

To date, in vivo evidence detailing immunological communication between the brain and skull bone marrow in humans remains limited. Through PET imaging, inflammation has been detected in the skull bone marrow adjacent to regions of cortical spreading depression in migraine patients ^6^. Additionally, Ringstad et al. recently used a magnetic resonance imaging (MRI) contrast agent as a CSF tracer, revealing molecular efflux from CSF to the skull bone marrow in individuals with CSF circulation disorders ^7^. Nevertheless, despite the noteworthy variations in the number, length, and diameter of SMCs being observed ^5^, the specific flow patterns of CSF in different skull bone marrow regions have not been fully characterized.

Recent research has highlighted the capacity of the skull bone marrow in contributing myelomonocytic cells to the surrounding meningeal tissue ^8^. In mice, Padi et al. demonstrated the migration of green fluorescent protein-positive streptococcus pneumoniae from CSF into the skull bone marrow during episodes of bacterial meningitis ^9^. However, similar observations in human subjects have yet to be documented, leaving the specific roles and influencing factors of these channels open to further exploration. Aging, combined with geriatric diseases such as hypertension and diabetes, increases susceptibility to systemic inflammation ^10^, which may be associated with CSF drainage in the skull bone marrow. Additionally, sleep disorders are reported to be related to systemic inflammation and could also impact CSF drainage ^11, 12^. Furthermore, since the impact of meningeal immunity and CSF drainage function has been reported to influence cognitive function and neurodegenerative diseases such as Alzheimer’s disease ^13, 14^, we thus hypothesize that CSF drainage in the skull bone marrow might affect cognitive function.

In this investigation, we utilized intrathecal contrast-enhanced MRI, incorporating three-dimensional T1-weighted (3D-T1) scans at multiple time points: baseline (prior to intrathecal administration of contrast agent), and subsequently 4.5 hours, 15 hours, and 39 hours post-administration in a cohort with neurological conditions. Gadolinium, utilized as a CSF tracer, was intrathecally administered to patients who were indicated for lumbar puncture. CSF drainage function was characterized as the percentage changes in the signal unit ratio within the skull bone marrow from baseline to time points post administration of gadolinium. We aimed to: 1) ascertain the ability of bone marrow within specific skull regions to facilitate CSF drainage; 2) examine the variables influencing the CSF drainage efficacy of the skull bone marrow; and 3) delineate the potential physiological and pathological functions associated with this drainage mechanism within the skull bone marrow.

## Materials and Methods

### Participants

The study enrolled patients with indications for lumbar puncture who voluntarily participated. Exclusion criteria comprised a history of known adverse reactions to contrast agents, severe allergic reactions in general, renal dysfunction, and pregnant or breastfeeding. Because this is a clinically-indicated, heterogeneous cohort without healthy volunteers, analyses are descriptive and association-focused, and findings are not intended to establish normative values or causal effects.

Prior to lumbar puncture, all patients underwent MRI sequences (baseline). Intrathecal injection of gadolinium was administered to all patients between 3:30 and 4:00 pm on day 1. Subsequent MRI sequences were then conducted at 4.5 hours (approximately 8:00∼8:30 pm on day 1), 15 hours (around 6:30∼7:00 am on day 2) and 39 hours (around 6:30∼7:00 am on day 3).

### Ethics statement

The study, incorporating the administration of intrathecal gadolinium agents, received approval from the Ethics Committee of Second Affiliated Hospital, School of Medicine, Zhejiang University (Approval Number: YAN-2018-111). All clinical investigations were conducted in compliance with the principles outlined in the Declaration of Helsinki. Written informed consent was obtained for all the participants.

### Intrathecal administration of gadolinium

The contrast agent was intrathecally injected at the L3-4 or L4-5 lumbar intervertebral space. The injection consisted of 1 mL of 0.5 mmol/mL gadolinium (Omniscan; GE Healthcare). Prior to injection, the correct positioning of the syringe tip in the subarachnoid space was confirmed by observing CSF backflow from the puncture needle. Following the intrathecal injection, patients were instructed to rotate themselves around the long axis of the body twice and then maintain a supine position for the next 4 hours.

### MRI protocol

A 3.0T MRI scanner (GE 750; GE Healthcare, Chicago, IL) was used, maintaining consistent imaging protocol settings across all time points. The imaging acquisitions comprised axial head 3-dimensional T1-weighted imaging (3D-T1). The principal imaging parameters included: repetition time = 7.3 ms, echo time = 3.0 ms, flip angle = 8°, thickness = 1 mm, field of view = 25 × 25 cm^2^, matrix = 250 × 250 pixels.

### Evaluation of imaging

Circular regions of interest were placed at predefined locations in the skull bone marrow, subarachnoid space and cortex [near the superior sagittal sinus (SSS), near the lateral fissure, and near the cisterna magna], as well as venous sinus and nasal turbinates on 3D-T1 images on baseline, and at 4.5 hours, 15 hours and 39 hours after the intrathecal administration of gadolinium. RadiAnt (Medixant, Poznan, Poland), a Digital Imaging and Communications in Medicine (DICOM) viewer, was utilized for this purpose. We recorded and normalized the mean signal unit of each region of interest against the reference. Based on previous studies, the vitreous body of the ocular bulb was chosen as the reference due to the absence of significant tracer accumulation following the intrathecal injection of gadodiamide ^15^. For each time point, the signal unit ratio between the regions of interest and the reference was determined. The percentage changes in the signal unit ratio from baseline to 4.5 hours, 15 hours, and 39 hours were calculated. For each patient, we compared the signal unit ratio at 4.5 hours, 15 hours, and 39 hours at each region. The time point exhibiting a relatively high signal unit ratio was defined as the peak time point of CSF tracer enrichment.

### Sleep quality and cognition assessment

The Pittsburgh Sleep Quality Index (PSQI), was employed to assess the quality and patterns of sleep, encompassing seven domains: subjective sleep quality, sleep latency, sleep duration, habitual sleep efficiency, sleep disturbances, use of sleeping medication, and daytime dysfunction. Participants were asked to report on their sleep experiences for past month preceding the initial MRI examination in the hospital. The total score ranged from 0 to 21, with higher scores indicating more severe dyssomnia ^16^. The Telephone Montreal Cognitive Assessment (T-MoCA) was used to evaluate the cognitive function, consisting of five domains: attention and calculation, language, abstraction, delayed recall and orientation. The assessment was conducted one month after discharge over the phone. The total score was 22 points, with higher scores indicating better cognitive function ^17^.

### Statistical analysis

Continuous variables were described as mean with standard deviations or medians with interquartile spacing, while categorical variables were presented as numbers and percentages. Differences between continuous or categorical data within the same individual were determined using paired-sample t tests or paired-sample Wilcoxon signed rank test, respectively. Correlations between continuous or rank data were determined using Spearman correlation analysis. The Benjamini–Hochberg method was used to correct for multiple comparisons. Multivariable analyses were conducted using linear regression analysis. Differences in continuous data were assessed using linear mixed models with a random intercept. Bootstrapping technique with 10,000 replicate samples with replacement was used to calculate the empirical power of linear mixed models.

Mediation analyses were used to assess whether the association between sleep (PSQI total scores) and cognitive function (T-MoCA total scores) might be mediated by the CSF drainage function of the skull bone marrow near the SSS at 4.5 hours. It involved testing three pathways: step 1, the association of sleep with cognitive function; step 2, the association of sleep with the CSF drainage function; and step 3, the association of the CSF drainage function of skull bone marrow near the SSS at 4.5 hours with cognitive function when controlling for sleep. Multivariable regression analysis was performed for each pathway, adjusting for age, education years, diagnosis of neurodegenerative disease, hypertension, and diabetes. If all associations were satisfied, the indirect effect was estimated. The extent of the indirect effect was estimated by examining the change in the β of PSQI total scores for T-MoCA total scores before and after including the CSF drainage function of the skull bone marrow near the SSS. The product of coefficient approach was also adopted for mediation analysis. The 95% confidence interval of the indirect effect was calculated using the bootstrapping technique with 5000 replicate samples with replacement. Statistical analyses were performed using SPSS software version 26.0 (IBM, Armonk, NY), and statistical significance was set at *p* = 0.05 (2-tailed).

## Results

### Study cohort

In the current study, a total of 87 patients were enrolled, with an average age of 58 ± 13 years. The patient diagnoses predominantly included peripheral neuropathy, encephalitis, and neurodegenerative diseases. Comorbidity analysis revealed diabetes in 31 patients (35.6%) and hypertension in 41 patients (47.1%). Detailed clinical data were summarized in **Table 1**. Cognitive function was assessed via the T-MoCA, with higher scores reflecting better cognitive performance, and the average score was 14.3 ± 4.1. Sleep quality was measured utilizing the PSQI, with lower scores indicating better sleep quality, and the average score was 6.6 ± 4.1.

**Table 1.** Clinical characteristics of all patients.

| Characteristics (n = 87) |  |
| --- | --- |
| Age (years) | 58 ± 13 |
| Female | 45 (51.7%) |
| Diabetes | 31 (35.6%) |
| Hypertension | 41 (47.1%) |
| Disease diagnosis |  |
| Peripheral neuropathy | 47 (54%) |
| Encephalitis | 15 (17.2%) |
| Normal pressure hydrocephalus | 4 (4.5%) |
| Suspected CSF leakage | 3 (3.4%) |
| Hepatic encephalopathy | 1 (1.1%) |
| Neurodegenerative diseases |  |
| Parkinson's disease | 5 (5.7%) |
| Motor neuron disease | 5 (5.7%) |
| Possible cerebral amyloid angiopathy | 3 (3.4%) |
| Multiple sclerosis | 1 (1.1%) |
| Multiple system atrophy | 1 (1.1%) |
| Alzheimer's disease | 1 (1.1%) |
| Lewy body dementia | 1 (1.1%) |
| Education years | 7 ± 5 |
| T-MoCA total scores (n = 58) | 14.3 ± 4.1 |
| Attention and calculation | 4.5 ± 1.6 |
| Language | 1.7 ± 1.0 |
| Abstraction | 0.6 ± 0.7 |
| Delayed recall | 2.1 ± 1.7 |
| Orientation | 5.3 ± 1.1 |
| PSQI total scores (n = 61) | 6.6 ± 4.0 |
| Sleep quality | 1.1 ± 0.9 |
| Sleep latency | 1.6 ± 1.1 |
| Sleep duration | 0.8 ± 0.9 |
| Habitual sleep efficiency | 0.7 ± 1.0 |
| Sleep disturbances | $1.0 \pm 0.5$ |
| Use of sleeping medication | $0.3 \pm 1.0$ |
| Daytime dysfunction | $1.1 \pm 1.2$ |

CSF: cerebrospinal fluid; T-MoCA: Telephone Montreal Cognitive Assessment; PSQI: Pittsburgh Sleep Quality Index.

### CSF tracer enrichment in skull bone marrow

On intrathecal contrast-enhanced MRI, we evaluated the signal unit ratio at each time point (baseline, 4.5, 15, and 39 hours) in three regions of the skull bone marrow, adjacent to the SSS, the lateral fissure, and the cisterna magna (**Figure 1**). We then calculated the percentage change of the signal unit ratio from baseline to 4.5, 15, and 39 hours in each region. Lower percentage change indicated better CSF drainage function. Post-gadolinium injection, significant alterations in the percentage changes were observed across all regions of skull bone marrow (all *p* < 0.05, **Figure 1p-r**), substantiating the migration of tracer from the CSF into the skull bone marrow. This tracer enrichment phenomenon was also evident in patients with peripheral neuropathy (all *p* < 0.05, **Supplemental Figure 1**). Specifically, near the SSS, the tracer reached its peak at 4.5 hours in 67.9% (55/81) of patients, at 15 hours for 18.5% (15/81), and at 39 hours for 2.5% (2/81), with 11.1% (9/81) showing no significant change. Near the lateral fissure, peak tracer enrichment at 4.5 hours was observed in 36.4% (27/74) of patients, at 15 hours in 32.4% (23/74), and at 39 hours in 21.6% (16/74), with 9.4% (7/74) displaying no notable change. Near the cisterna magna, the peak was at 4.5 hours for 18.0% (13/72) of patients, at 15 hours for 50.0% (36/72), and at 39 hours for 23.6% (17/72), with the remaining 8.3% (6/72) exhibiting no significant shift.

**Figure 1:**
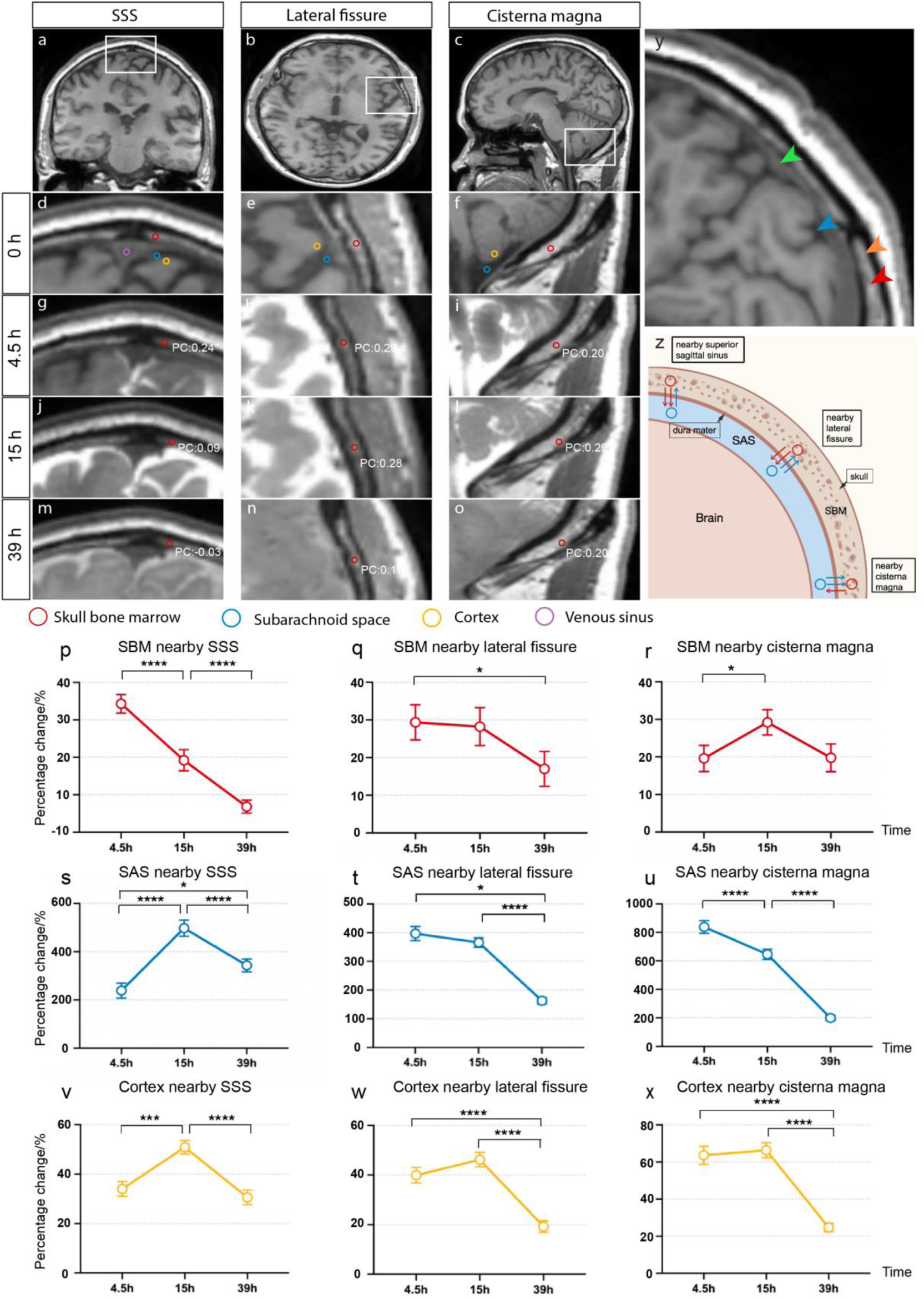
Dynamic patterns of cerebrospinal fluid (CSF) drainage and signal changes across brain regions following intrathecal contrast administration **a-c** Three anatomical planes for region of interest (ROI) placement. ROIs were placed in the skull bone marrow (SBM) (red circle), subarachnoid space (SAS) (blue circle) and cortex (yellow circle) adjacent to three anatomical landmarks: the superior sagittal sinus (SSS) (first column), lateral fissure (second column) and cisterna magna (third column). The venous sinus is indicated by the lavender circle. White boxes in the top row images indicate regions that are magnified in subsequent rows. **d-o** Sequential MRI images show ROIs at different time points following intrathecal gadodiamide administration: baseline (0 h), 4.5 hours (4.5 h), 15 hours (15 h), and 39 hours (39 h). Percentage change (PC) values are labeled to illustrate signal variations over time. **p-x** Percentage change in signal unit ratio from baseline to post-contrast administration in the SBM (**p-r**), SAS (**s-u**), and cortex (**v-x**) adjacent to the SSS, lateral fissure and cisterna magna. Data are presented as mean ± standard error of the mean. Statistically significant differences between time points, identified using linear mixed model analysis, are indicated by asterisks: * *p* < 0.05, *** *p* < 0.001, **** *p* < 0.0001. **y** High-magnification image identifying key structures with arrowheads: skull (orange), SBM (red), SAS (blue), and dura mater (green). **z** Hypothesized patterns of SBM drainage. There is potential bidirectional communication between CSF in the SBM and SAS. Drainage in SBM near the SSS is likely relatively rapid, mainly flowing into the SAS. Conversely, near the cisterna magna, CSF in the SAS primarily drains into the SBM. Intermediate regions, such as those near the lateral fissure, may exhibit characteristics of balanced bidirectional drainage. Given the unbalanced drainage near the SSS and cisterna magna, we hypothesize that there might be an alternative drainage pathway in between skull and brain, which could expedite the flow of CSF from areas neighboring the SBM adjacent to the cisterna magna to those near the SSS. Moreover, the early peak enrichment at 4.5 hours in SBM suggests that this pathway may be very rapid.

To investigate potential communication between CSF in the subarachnoid space and the skull bone marrow, we evaluated the percentage changes of CSF in the subarachnoid space near the regions mentioned before (**Figure 1s-u**). Remarkably, the average percentage changes in the skull bone marrow near the SSS reached peak at 4.5 hours, earlier than that observed within the subarachnoid space nearby (**Figure 1p,s**), suggesting the drainage of CSF tracer through the skull bone marrow might be a notably fast route.

The observed CSF drainage patterns within the skull bone marrow and subarachnoid space near the lateral fissure were aligned, with the percentage changes both peaking at 4.5 hours (**Figure 1q,t**). The peak time point for the skull bone marrow near the cisterna magna was recorded at 15 hours, later than the peak time observed in the surrounding subarachnoid space (**Figure 1r,u**). Given the unbalanced drainage near the SSS and cisterna magna, we hypothesize that there might be an alternative drainage pathway in between skull and brain, which could expedite the flow of CSF from areas neighboring the skull bone marrow adjacent to the cisterna magna to those near the SSS. In addition, positive correlations were noted in the percentage changes between the bone marrow and subarachnoid space near the lateral fissure at both 4.5 and 39 hours, as well as the percentage change between the skull bone marrow and subarachnoid space near the cisterna magna at 4.5 hours, 15 hours, and 39 hours post-contrast (**Supplemental Figure 2**). These results indicated the potential drainage of CSF tracer directly into the skull bone marrow near both the lateral fissure and cisterna magna. **Figure 1z** depicted hypothesized mechanism of skull bone marrow drainage patterns.

To assess the potential contribution of venous signals to skull bone marrow tracer enrichment, we analyzed the signal changes in the venous sinus (**Supplemental Figure 3a**). Additionally, we observed correlations between skull bone marrow and venous sinus signals at several time points, indicating potential venous involvement (**Supplemental Figure 4**). We also assessed tracer enrichment in the cortex (**Figure 1v-x**) and its relationship with skull bone marrow tracer enrichment. While some correlations were observed, they were generally weaker compared to those with the subarachnoid space, suggesting a limited direct interaction between skull bone marrow drainage and cortical tracer dynamics (**Supplemental Figure 5**).

To explore the relevance of olfactory CSF efflux in humans, we conducted an additional analysis focusing on regions adjacent to the nasal turbinates, which are anatomically linked to olfactory CSF drainage pathways. The analysis revealed a gradual increase in the PC of the contrast agent signal from 4.5 hours to 15 and 39 hours post-intrathecal administration, with statistically significant differences between these time points (*p* < 0.05) (**Supplemental Figure 3b-d**). This delayed enrichment pattern contrasts with the more rapid tracer accumulation observed in the skull bone marrow regions, suggesting that olfactory CSF efflux is not a rapid drainage pathway in humans.

### Influencing factors for CSF drainage function of skull bone marrow

Given the fastest CSF drainage observed, we focused on the CSF drainage function of the skull bone marrow adjacent to the SSS, which was defined as the percentage change in signal unit ratio from baseline to 4.5 hours, where a lower percentage change indicates superior drainage function. We found that impaired drainage function was significantly correlated with female sex, hypertension, diabetes, and aging (all *p* < 0.05, **Figure 2a-d**). Additionally, a positive correlation was observed between drainage function and neutrophil percentage (*p* = 0.032, **Figure 2g**), while negative correlations were noted with lymphocyte (*p* = 0.035, **Figure 2h**) and monocyte percentages (*p* = 0.02, **Figure 2i**), supporting an association between drainage and systemic inflammation. After adjusting for age, the monocyte percentage remained significantly associated with the drainage function (β = -0.278, *p* = 0.008). In peripheral neuropathy subgroup (n = 45), impaired drainage function was linked to hypertension (*p* = 0.008) and diabetes (*p* = 0.007), but not to sex (*p* = 0.49) or aging (*p* = 0.06, **Supplemental Figure 6a-d**). Remarkably, after adjusting for hypertension, age, and sex, diabetes continued to show a significant association with impaired drainage function (β = 0.338, *p* = 0.03).

**Figure 2:**
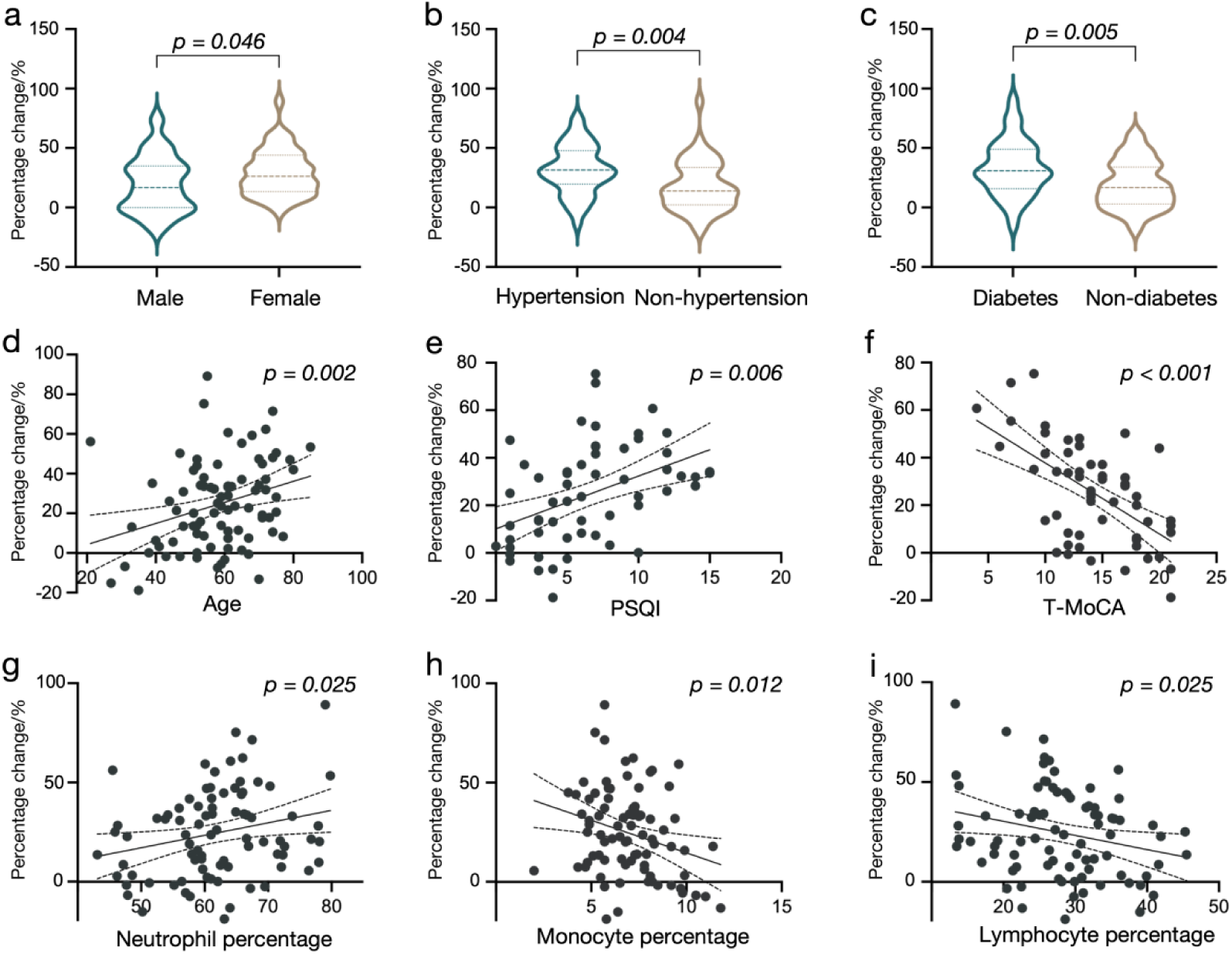
Influencing factors related to drainage function in skull bone marrow (SBM) near the superior sagittal sinus (SSS). Female (**a**), patients with hypertension (**b**) or diabetes mellitus (**c**) exhibit lower percentage changes of signal unit ratio from baseline to 4.5 hours in SBM near the SSS, compared to male and patients without these conditions. Age (**d**) and the Pittsburgh Sleep Quality Index (PSQI) total scores (**e**) are positively correlated with the percentage change, while the Telephone Montreal Cognitive Assessment (T-MoCA) total scores (**f**) was negatively correlated with the percentage change. Neutrophil percentage (**g**) is positively related, while monocyte (**h**) and lymphocyte percentages (**i**) are negatively correlated with the percentage change. Each plot displays the sample size, the fitted regression line, and the respective Spearman or Pearson correlation coefficients (ρ or R), with corresponding *P*-values.

### Relationship between CSF drainage function in skull bone marrow and sleep quality

The drainage function of the bone marrow near the SSS showed negative correlations with the PSQI total score, along with sub-items including subjective sleep quality, sleep latency, sleep disturbances, and daytime dysfunction (all FDR *p* < 0.05, **Table 2**). This association persisted even after adjusting for age, sex, and neurodegenerative diseases, indicating an independent relationship between PSQI total score and the drainage function near the SSS (β = 0.373, *p* = 0.001). To mitigate the impact of intracranial diseases on sleep, a peripheral neuropathy subgroup analysis was undertaken (n = 34, **Supplemental Figure 6e**). This analysis uncovered a negative correlation between the drainage function and the PSQI total scores (*p* = 0.013), further emphasizing the relationship between sleep quality disturbances and compromised cranial marrow drainage.

**Table 2.** Correlations between sleep quality and its sub-items and percentage change of signal unit ratio from baseline to 4.5 hours in SBM near the SSS.

|  | PSQI<br>total<br>scores | Sleep<br>quality | Sleep<br>latency | Sleep<br>duratio<br>n | Habitu<br>al<br>sleep<br>efficie<br>ncy | Sleep<br>disturb<br>ances | Use of<br>sleepin<br>g<br>medica<br>tion | Dayti<br>me<br>dysfun<br>ction |
| --- | --- | --- | --- | --- | --- | --- | --- | --- |
| $\rho$ | 0.390 | 0.297 | 0.380 | 0.090 | 0.128 | 0.263 | 0.200 | 0.328 |
| FDR $P$ | 0.012 | 0.045 | 0.012 | 0.499 | 0.381 | 0.070 | 0.171 | 0.0293 |

SBM: skull bone marrow; SSS: superior sagittal sinus; PSQI: Pittsburgh Sleep Quality Index; All $P$ value were corrected false discovery rate by the Benjamini-Hochberg method in PSQI total scores and each domain respectively.

### Relationship between CSF drainage function of skull bone marrow and cognitive function

The drainage function of the bone marrow adjacent to the SSS showed a positive correlation with the total scores of the T-MoCA, as well as specific domains such as attention and calculation, delayed recall, and orientation (all FDR *p* < 0.05, **Table 3**). After controlling for confounding factors including age, sex, years of education, and the presence of neurodegenerative diseases, this drainage function was identified as an independent protective factor for T-MoCA total scores (β = -0.381, *p* = 0.002). In the subgroup of individuals with peripheral neuropathy, a positive relationship was also observed between the drainage function and T-MoCA total scores (n = 34, *p* = 0.011, **Supplemental Figure 6f**).

**Table 3.**
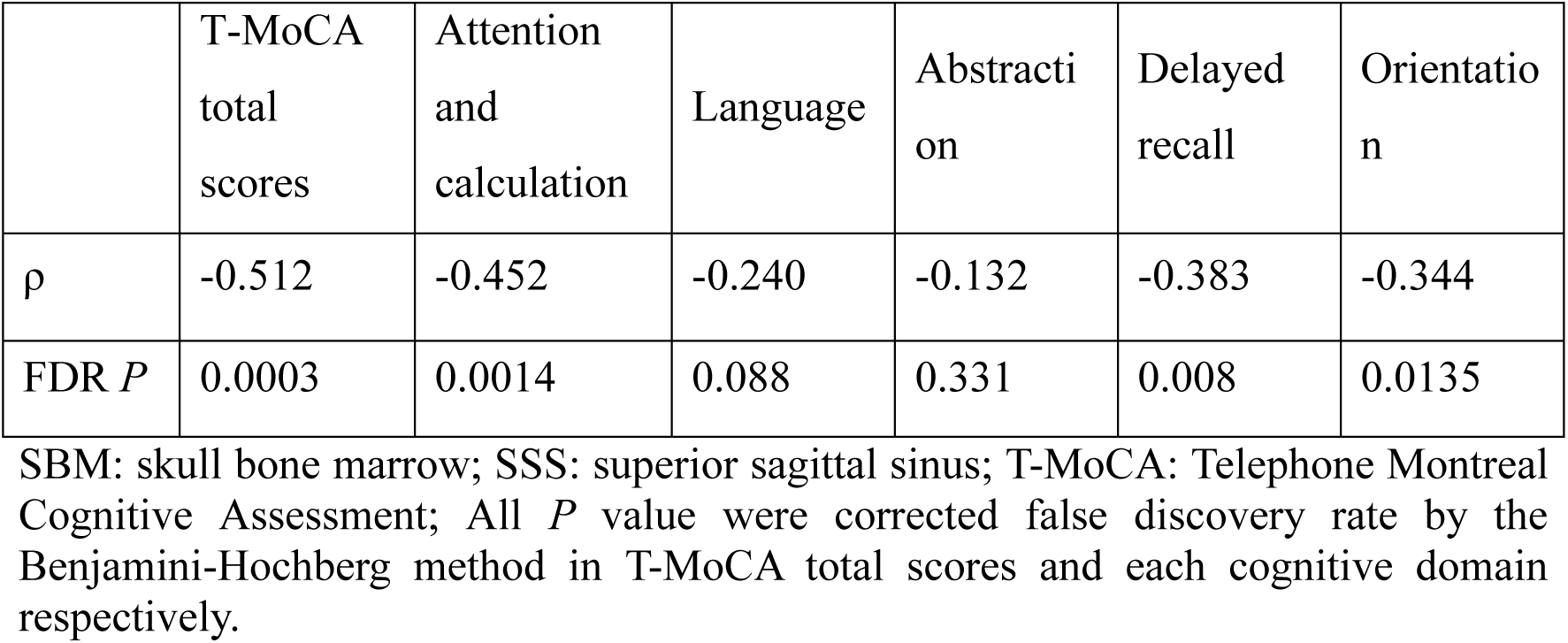
Correlations between cognitive function and its sub-items and percentage change of signal unit ratio from baseline to 4.5 hours in SMB near the SSS.

### CSF drainage function of skull bone marrow served as a mediator in the association between sleep quality and cognitive decline

Among 56 participants who completed intrathecal contrast-enhanced MRI alongside assessments of sleep and cognitive functions, we explored the mediation role of CSF drainage through skull bone marrow in association between sleep quality and cognitive performance. Comprehensive clinical details of these subjects were provided in **Supplemental Table 1**. When age, education years, the presence of neurodegenerative diseases, hypertension, and diabetes were considered as covariates, the effect size of PSQI total scores on T-MoCA total scores significantly decreased after controlling the drainage function of the bone marrow near the SSS (change in β [bootstrap 95% confidence interval]: -0.092 [-0.195, -0.004]; **Figure 3**). Thus, the CSF drainage function, serves as a mediator, explaining 38% of the association between PSQI total scores and T-MoCA total scores.

**Figure 3:**
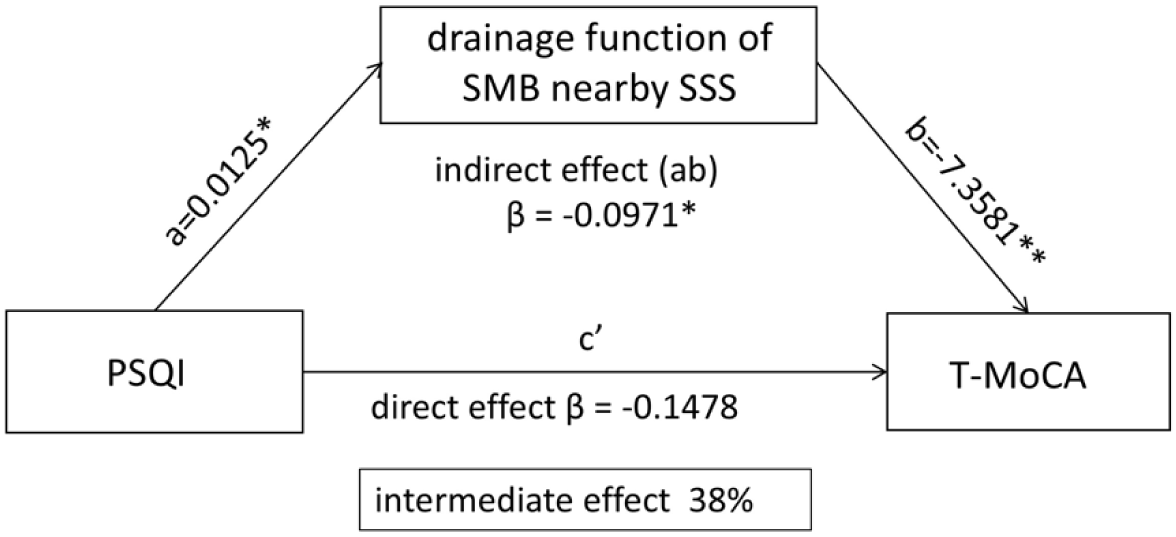
Mediation analysis evaluating whether the association of sleep quality with cognitive performance is mediated by the drainage function of the skull bone marrow (SBM) adjacent to the superior sagittal sinus (SSS). The direct effect of Pittsburgh Sleep Quality Index (PSQI) score on Telephone Montreal Cognitive Assessment (T-MoCA) score is denoted by c’ (β = -0.1478). The indirect effect through the drainage function of SBM adjacent to the SSS is captured by path a (β = 0.0125*) to path b (β = -7.3881**), resulting in an indirect effect (ab) of β = -0.0971*. The intermediate effect explains 38% of the total effect, highlighting the significant mediating influence of CSF drainage efficiency on cognitive outcomes relative to sleep quality. Asterisks denote levels of statistical significance with covariates considered: * *p* < 0.05, ** *p* < 0.01.

## Discussion

In the current study, we observed tracer signal changes consistent with CSF drainage into the skull bone marrow, highlighting the drainage variations nearby different regions, based on a cohort of patients who underwent intrathecal contrast-enhanced MRI. Our findings indicate that the drainage in the skull bone marrow adjacent to the SSS may represent the most rapid pathway. This process is significantly influenced by age, underscoring the pivotal role of aging in this physiological mechanism, with comorbidities such as hypertension and diabetes also exerting notable effects. The association of inflammatory cells with CSF drainage suggests that systemic inflammation significantly impacts the capacity for drainage. Importantly, this drainage function explains 38% of the correlation between PSQI total scores and T-MoCA total scores, indicating a substantial role in cognitive decline associated with sleep disturbances.

Researches over the years has increasingly focused on the connection between the brain and the skull bone marrow. In 2018, Fanny H et al. uncovered direct vascular channels connecting the skull bone marrow with the surface of the CNS in rodents by confocal microscopy ^4^. The discovery was further supported by studies that demonstrated the movement of CSF tracers from the cisterna magna into the skull bone marrow through these channels, thereby delineating a tangible route for CSF and bone marrow intercommunication ^9^. In 2022, Ringstad et al. reported CSF tracer enrichment within diploe of skull bone marrow in human participants with suspicion of CSF disorders by MRI ^7^. Not confined to the individuals with CSF disorders, our study highlights that CSF efflux into the skull bone marrow could be a broader phenomenon.

Importantly, we observed a rapid peak in tracer enrichment, compatible with earlier apparent dynamics near the SSS. This phenomenon may be attributed to the anatomical proximity of the parietal lobe to the SSS. In human studies, the most permeable channels to the sub-dural space are typically located beneath the arachnoid granulation, where they often penetrate the dura mater ^18^. These channels, with widths ranging from 40 to 90 micrometers, are frequently adjacent to larger blood vessels, which exceed 150 micrometers in width. The parietal region is noted to have a greater abundance of these channels compared to the frontal and temporal regions ^5^. Furthermore, CSF is known to drain from the subarachnoid space into the venous system, particularly through the superior sagittal sinus. Consequently, CSF in the vicinity of the superior sagittal sinus is more likely to flow from the skull bone marrow to the subarachnoid space. Moreover, recent anatomical studies involving cadaver dissections have unveiled the CSF canalicular system, a network of channels that flank the SSS and communicate with subarachnoid CSF through Virchow-Robin spaces ^19^.

Furthermore, we observed the unbalanced drainage near the SSS and cisterna magna. Consequently, we hypothesize that there might be an alternative drainage pathway in between skull and brain, which could expedite the flow of CSF from areas neighboring the skull bone marrow adjacent to the cisterna magna to those near the SSS. Recently, Møllgård et al. demonstrated the existence of a fourth meningeal layer in mice and human brains. This layer compartmentalizes the subarachnoid space and is designated the subarachnoid lymphatic-like membrane (SLYM) ^20^. Plá et al. have speculated that this arrangement allows fresh CSF, produced in the ventricles, to be directed preferentially upwards along the cerebral arteries, rather than mixing with the larger pool of CSF surrounding the brain ^21^. More recently, Smyth et al. identified discontinuities in the arachnoid barrier around bridging veins, termed arachnoid cuff exit (ACE) points. These ACE points permit the exchange of tracers between the dura mater and the subarachnoid space, enabling the clearance of CSF tracers. They also proposed a working model for CSF efflux, in which CSF accesses the dura mater directly through bulk flow before being drained by conventional lymphatics ^22^. The complex stratification of the meninges and the direction of fluid flow may explain why some CSF rapidly moves to other locations near the skull’s bone marrow. However, since transcranial imaging cannot distinguish the compartments, this hypothesis awaits confirmation in future studies.

We further revealed that impaired drainage function of skull bone marrow was influenced by aging, hypertension, and diabetes, identifying these factors as being associated with less favorable drainage. It may due to changes in skull structure and the abnormal expansion of marrow adipose tissue under the above conditions. Specifically, these factors are known to increase the risk of osteoporosis, which in turn affect bone architecture ^23, 24^. Additionally, researches have demonstrated that these elements significantly impede CSF circulation ^25-28^, ultimately affecting CSF drainage within the skull bone marrow.

Intriguingly, our analysis revealed that systemic inflammation, as indicated by the proportions of neutrophils, lymphocytes, and monocytes, significantly affects CSF drainage, highlighting an interaction between systemic health and neurophysiological processes. Actually, compared to subjects in previous research subjects ^7^, participants in our study are older and more frequently suffer from conditions like hypertension, diabetes, or degenerative diseases, therefore are more susceptible to chronic inflammatory conditions, which might be the cause of CSF efflux into the skull bone marrow. Furthermore, clinical studies have linked osteoporotic conditions, commonly observed in aging individuals and postmenopausal women, to a persistent, low-grade inflammatory environment characterized by changes in cytokine levels and the composition of immune cells ^29^. These findings underscore the potential relationship between inflammation, bone health, and CNS functionality, suggesting that systemic health could profoundly influence neurological functions by modulating CSF drainage pathways.

Our research highlights associations between CSF drainage function and sleep quality and cognitive performance. We observed a marked deceleration in CSF drainage within the bone marrow adjacent to the SSS in subjects with suboptimal sleep quality. Poor sleep quality has been found to increase amyloid β accumulation and promote neuroinflammation, particularly in hippocampal regions, leading to cognitive impairment ^12, 30^. Our findings provide evidence that the CSF drainage function of the skull bone marrow may be a crucial pathway to clear abnormal proteins and inflammatory molecules in patients with poor sleep. Notably, chronic sleep deprivation is associated with sustained systemic low-grade inflammation ^31^. In a meta-analysis of about 50,000 adults, sleep disturbance was associated with higher levels of C-reactive protein and interleukin-6 ^32^. Thus, we hypothesize that insufficient sleep may amplify chronic inflammation, thereby impeding drainage from the skull bone marrow and ultimately leading to cognitive deficits.

Based on these observations, we hypothesize that aging, hypertension, and diabetes induce structural and vascular changes in the skull bone marrow, directly impairing CSF drainage. These conditions also promote chronic systemic inflammation, which affects the permeability and function of meningeal and skull-marrow drainage pathways. Poor sleep quality acts as both a cause and consequence of impaired CSF clearance, creating a vicious cycle where sleep disturbances exacerbate inflammation, further hindering drainage and accelerating cognitive decline. This integrated framework suggests that CSF drainage dysfunction is not an isolated process but is tightly linked to systemic health, vascular integrity, immune regulation, and sleep physiology. Our findings underscore the potential of targeting CSF clearance pathways as a novel therapeutic approach to mitigate cognitive impairment and promote brain health.

Our study acknowledges several limitations that warrant attention. First, cohort heterogeneity and clinically indicated enrollment limit generalizability. Although we conducted subgroup analyses with a specific focus on peripheral neuropathy, caution should be considered when extrapolating our findings. Second, the manual placement of regions of interest could introduce potential measurement bias due to subjective judgment. While efforts were made to avoid cortical bone and minimize partial volume effects, some inherent measurement discrepancies may still remain. Third, the evaluation of sleep quality and cognitive function was based upon self-reported instruments. Future studies should incorporate more clinically robust diagnostic evaluation to substantiate our findings with greater precision. Fourth, our data collection was limited to three specific time points post-intrathecal injection, which may have missed significant fluctuations in CSF drainage patterns. Additionally, we were unable to precisely quantify the relative contributions of CSF, venous sources, and cortical tissue to the observed skull bone marrow tracer enrichment. While our analysis suggests that direct CSF drainage may predominate, correlations with venous sinus and cortical signals at certain time points indicate potential contributions from these compartments. Future studies with more frequent data points and advanced imaging techniques, such as dynamic contrast-enhanced MRI, tracer kinetics modeling, and BBB permeability assessments, will be essential to better characterize CSF drainage kinetics and clarify the extent of venous and cortical contributions. Fifth, despite these findings, our study is primarily descriptive, and the underlying biological mechanisms driving the observed CSF drainage alterations remain speculative. Although we propose potential pathways involving systemic inflammation, vascular remodeling, and metabolic effects on skull bone marrow structure, we lack direct mechanistic evidence confirming these relationships. To address this gap, we suggest that future studies employ advanced imaging techniques (e.g., dynamic PET imaging or high-resolution MRI of meningeal lymphatics) and incorporate biomarker analyses (e.g., inflammatory cytokines, bone marrow-derived immune cell profiling) to further elucidate the physiological and pathological mechanisms underlying skull bone marrow CSF drainage.

In summary, our study identified different CSF drainage patterns in the skull bone marrow and revealed their modulation by factors such as age, sex, and health conditions like hypertension and diabetes. We also found that systemic inflammation can impact the drainage, emphasizing the significance of CSF drainage intermediating sleep and cognitive health. This research advances our understanding of skull bone marrow neurophysiology and cognitive function mechanisms. Future research might explore targeted interventions to modulate CSF drainage in the skull bone marrow as a potential therapeutic strategy.

## Acknowledgement

We thank all participants and clinical staff involved in the study for their valuable contributions. This work was supported by grants from the National Natural Science Foundation of China (No. 81971101, 82171276, U23A20426).

## Author Contributions

ML and YZ contributed to the conception and design of the study. YZ, HJ, XZ, YL, ZZ, XH, HK, MF, and JS contributed to the acquisition and analysis of data. YZ and HJ drafted the text and prepared the figures.

## Data Availability Statement

Data presented in this article may be shared upon reasonable request to the corresponding or senior author by qualified investigators for non-commercial research use, subject to restrictions imposed by participant consent and applicable data protection regulations.

---

**Supplemental Figure 1:**
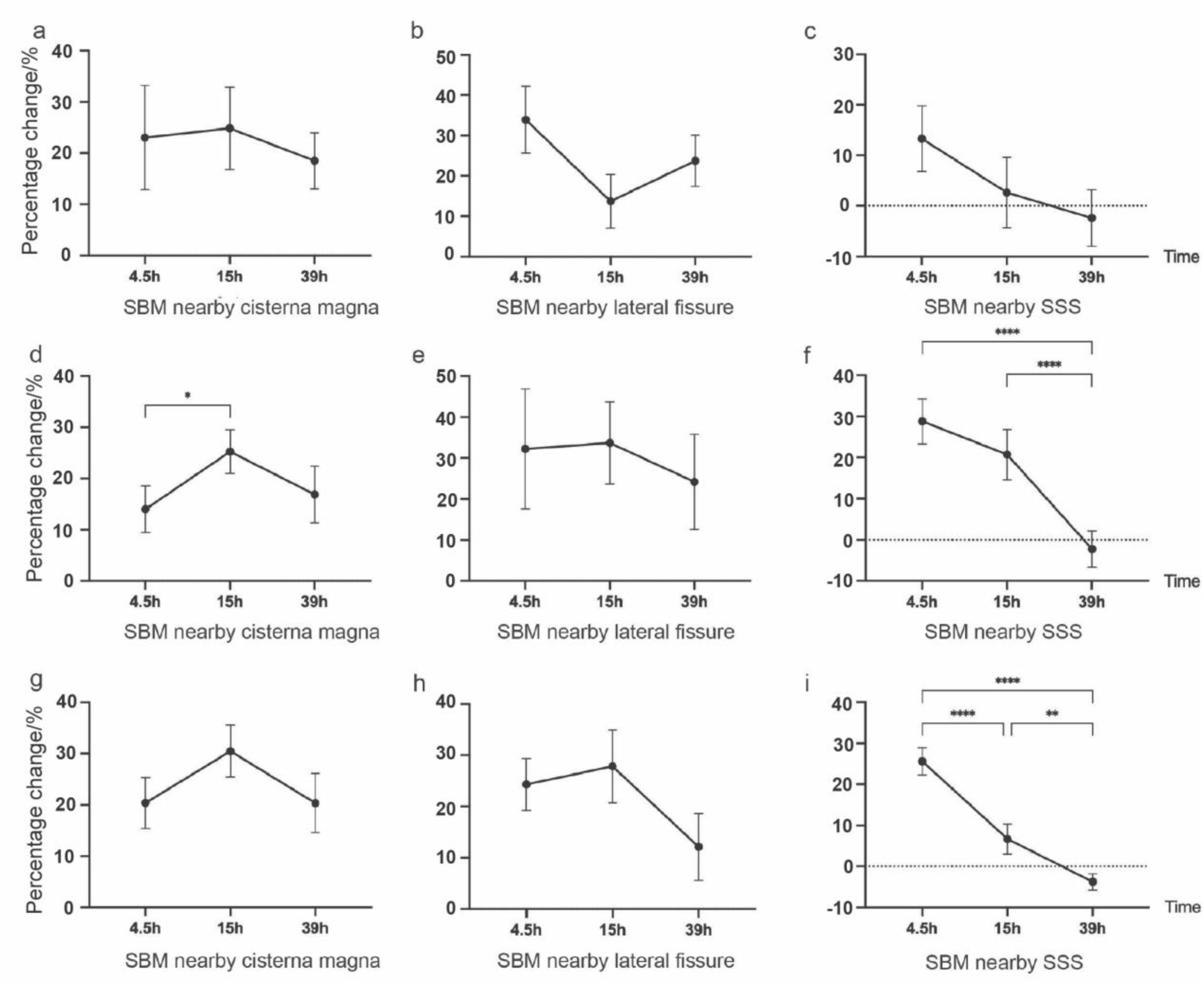
Tracer enrichment in skull bone marrow (SBM) among different diseases. This figure presents the percentage changes in the signal unit ratio from baseline to 4.5, 15, and 39 hours post cerebrospinal fluid (CSF) tracer administration across different SBM regions in patients with encephalitis (**a-c**, n = 15), neurodegenerative disease (**d-f**, n = 19), and peripheral neuropathy (**g-i**, n = 45). Error bars denote the standard error of the mean. Statistically significant differences between pairs of time points, as identified through linear mixed model analysis, are marked with stars: * *p* < 0.05, ** *p* < 0.01, **** *p* < 0.0001.

**Supplemental Figure 2:**
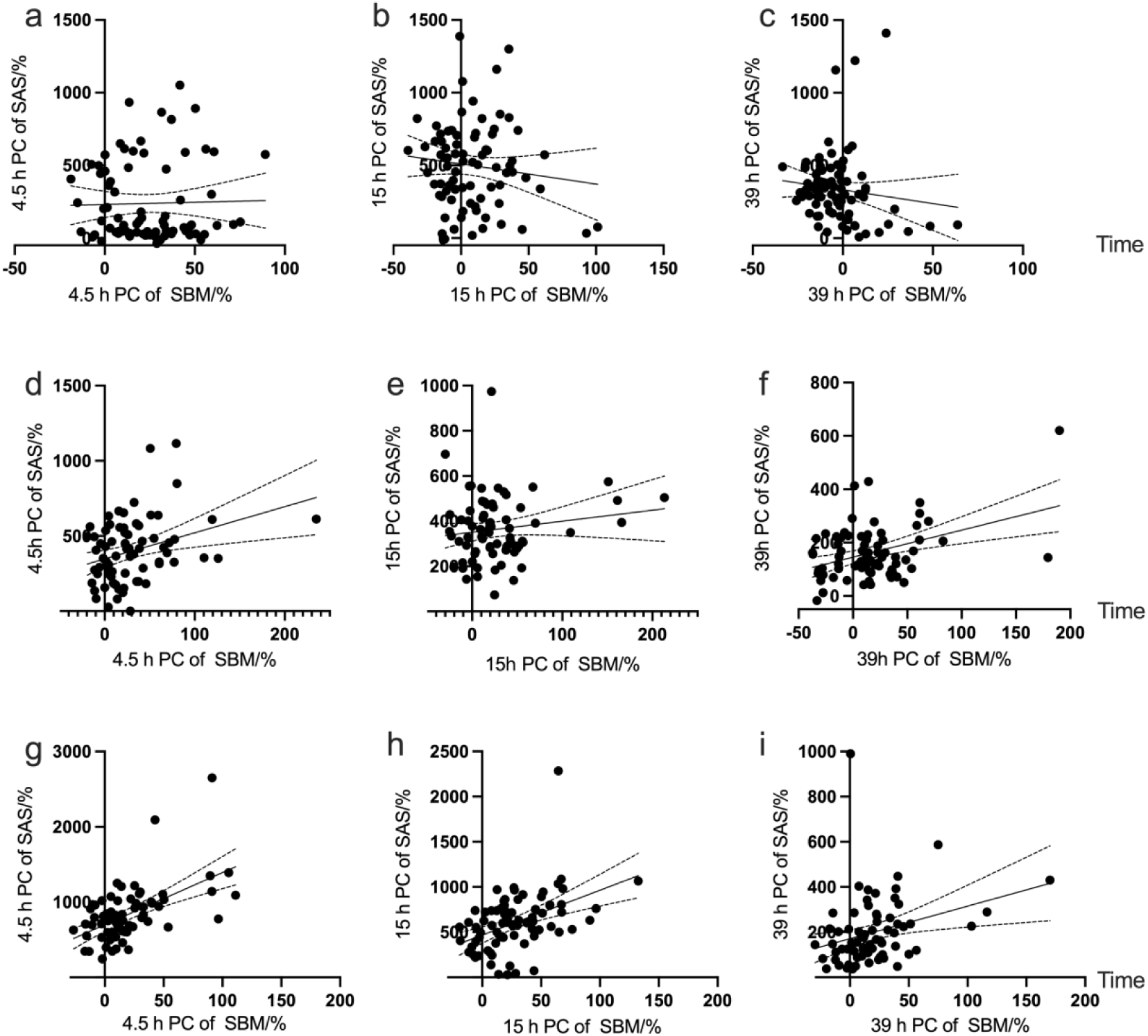
Correlation between tracer enrichment in skull bone marrow (SBM) and subarachnoid space (SAS). This figure elucidates the relationship between the percentage changes (PC) in the signal unit ratio of SBM and SAS from baseline to 4.5, 15, and 39 hours post cerebrospinal fluid (CSF) tracer administration in three anatomical regions: the superior sagittal sinus (**a-c**), the lateral fissure (**d-f**), and the cisterna magna (**g-i**). Each panel displays the sample size, a fitted regression line, and the Spearman correlation coefficient (ρ) alongside the associated *P*-value.

**Supplemental Figure 3:**
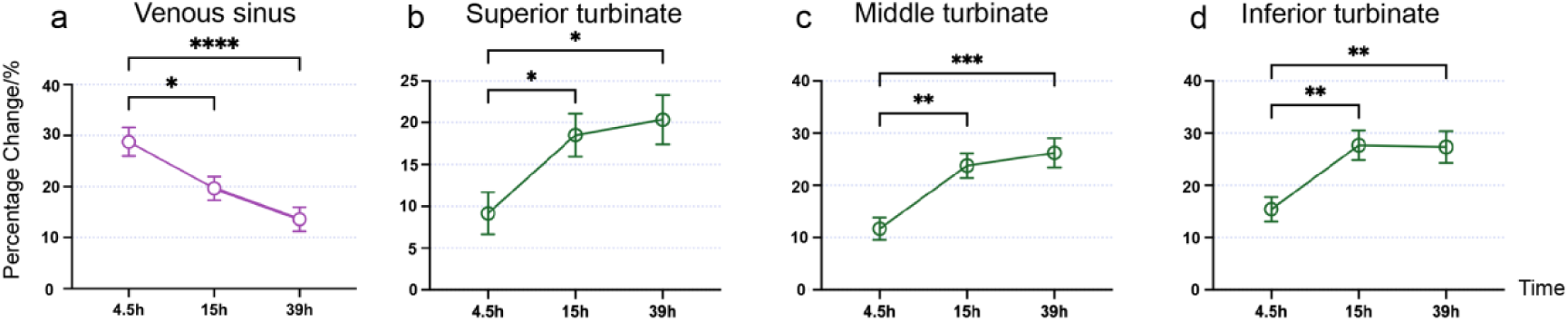
Tracer enrichment in venous sinus and nasal turbinates. **a** Percentage change in the signal unit ratio from baseline to 4.5, 15, and 39 hours post-cerebrospinal fluid (CSF) tracer administration in venous sinus. **b-d** Percentage change in the signal unit ratio from baseline to 4.5, 15, and 39 hours post-CSF tracer administration in the superior turbinate (**b**), middle turbinate (**c**), and inferior turbinate (**d**). Data are presented as mean ± standard error of the mean. Statistically significant differences between time points, identified using linear mixed model analysis, are indicated by asterisks: * *p* < 0.05, ** *p* < 0.01, *** *p* < 0.001, **** *p* < 0.0001.

**Supplemental Figure 4:**
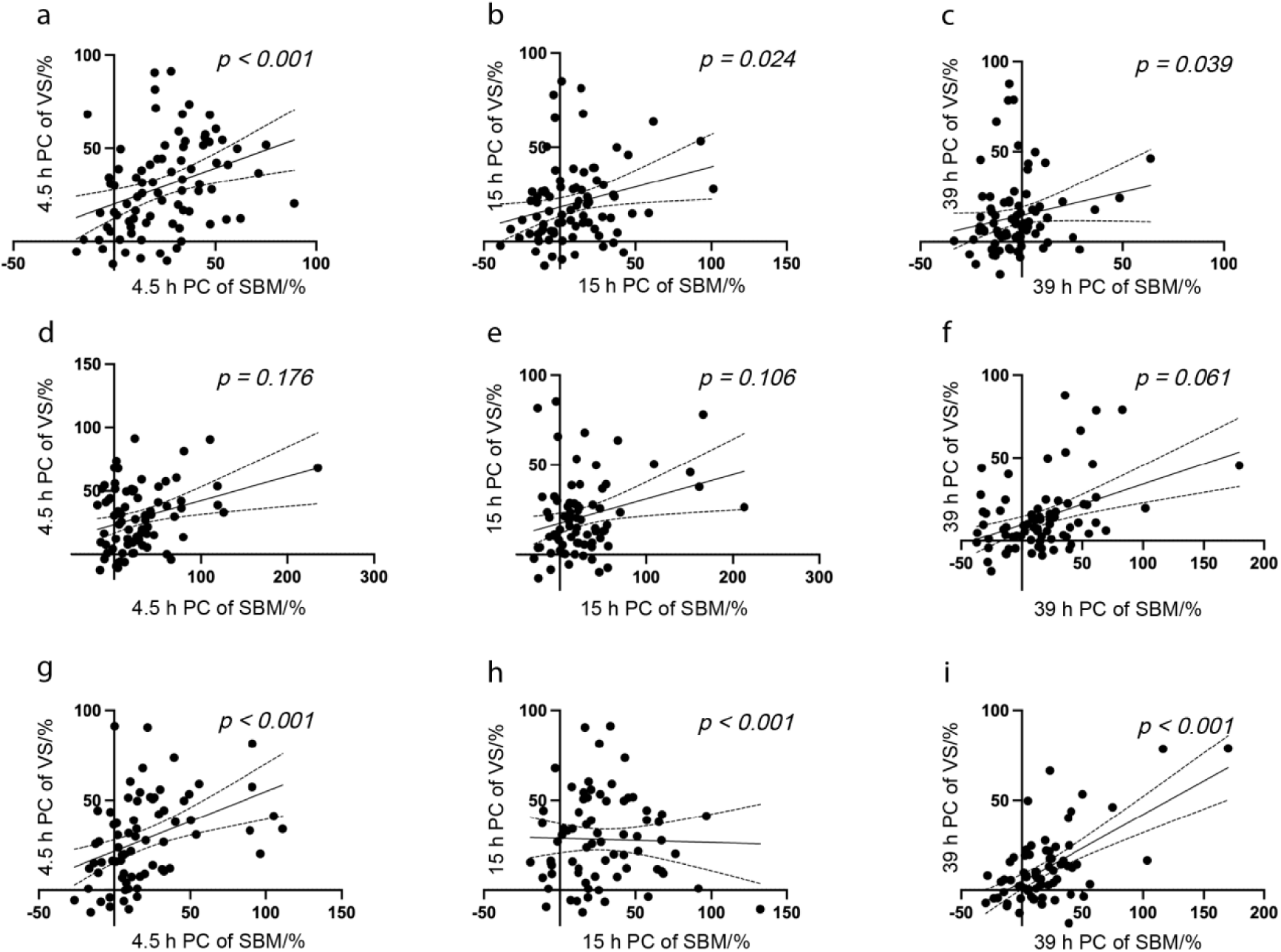
Correlation between tracer enrichment in skull bone marrow (SBM) and venous sinus (VS). This figure illustrates the relationship between the percentage changes (PC) in the signal unit ratio of SBM and the VS from baseline to 4.5, 15, and 39 hours post-cerebrospinal fluid (CSF) tracer administration in three anatomical regions: the superior sagittal sinus (SSS) (**a-c**), the lateral fissure (**d-f**), and the cisterna magna (**g-i**). Each panel displays the sample size, a fitted regression line with 95% confidence intervals (dotted lines), and the Spearman correlation coefficient (ρ) alongside the corresponding P-value, indicating the strength and statistical significance of the correlations observed.

**Supplemental Figure 5:**
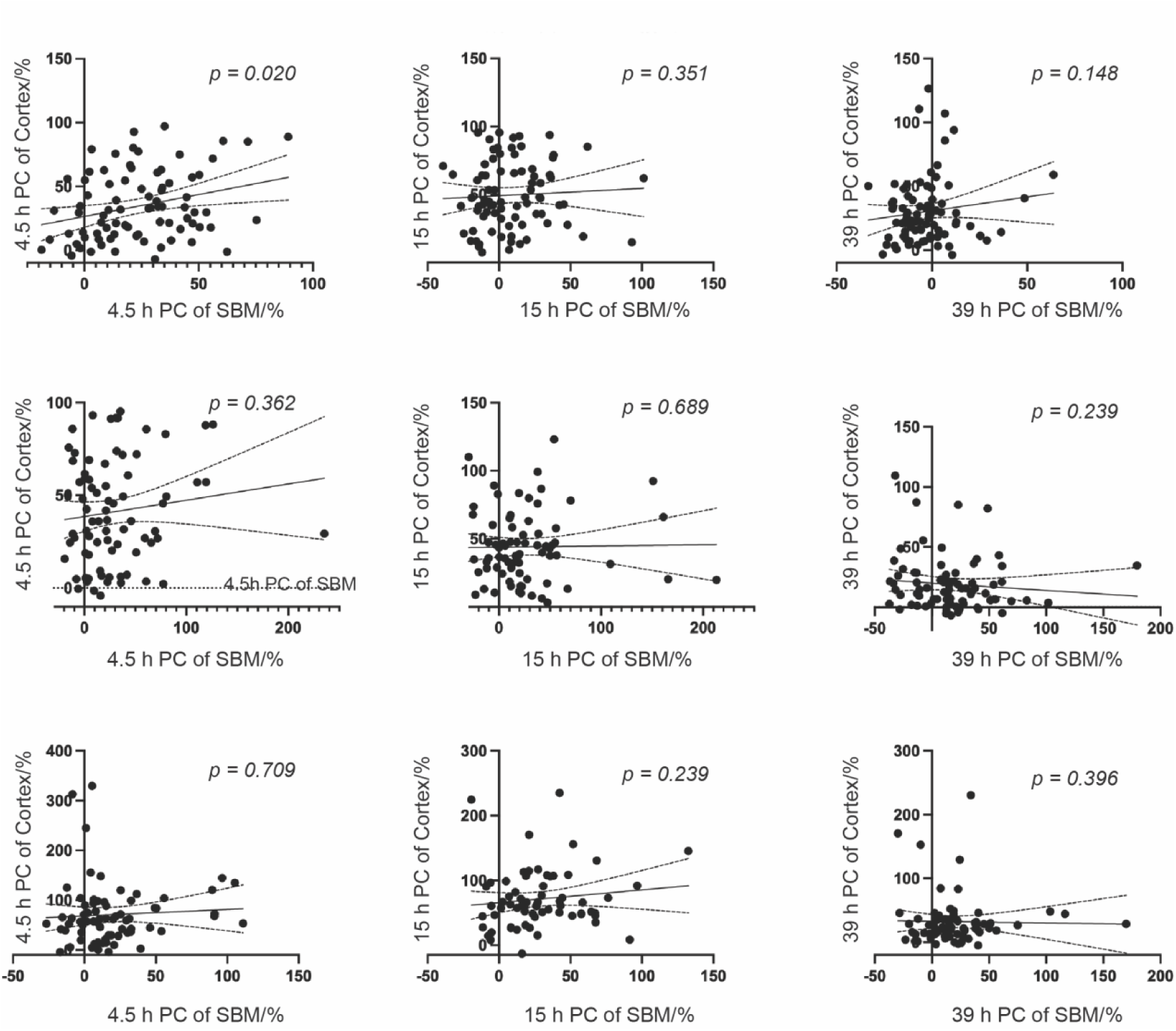
Correlation between tracer enrichment in skull bone marrow (SBM) and cortex. This figure illustrates the relationship between the percentage changes (PC) in the signal unit ratio of SBM and the cortex from baseline to 4.5, 15, and 39 hours post-cerebrospinal fluid (CSF) tracer administration in three anatomical regions: the superior sagittal sinus (SSS) (**a-c**), the lateral fissure (**d-f**), and the cisterna magna (**g-i**). Each panel displays the sample size, a fitted regression line with 95% confidence intervals (dotted lines), and the Spearman correlation coefficient (ρ) alongside the corresponding P-value, indicating the strength and statistical significance of the observed correlations.

**Supplemental Figure 6:**
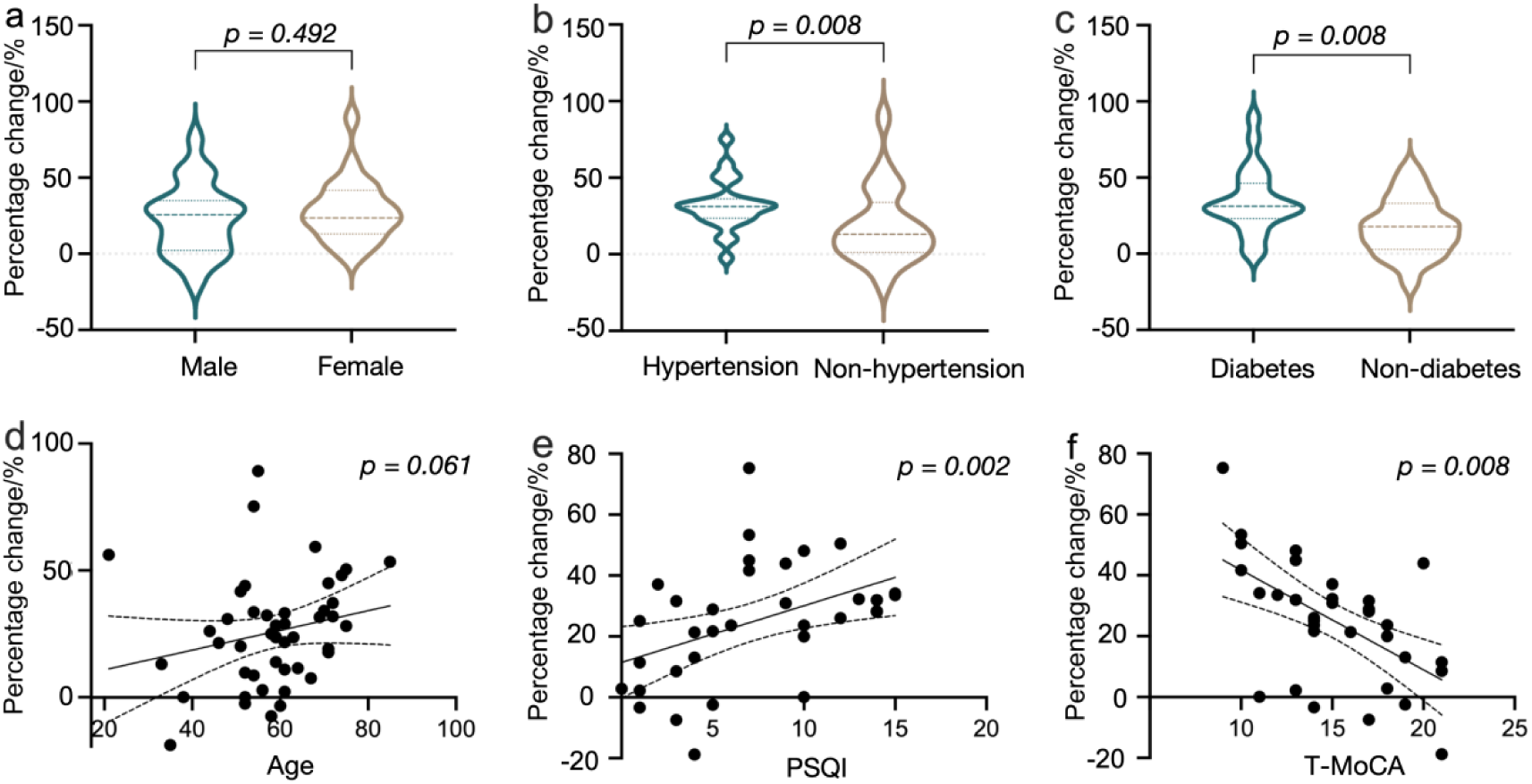
Influencing factors related to drainage function in SMB near the SSS in the subgroup analysis of patients with peripheral neuropathy. In the subgroup analysis of patients with peripheral neuropathy (n=45), no difference of the percentage changes of signal unit ratio from baseline to 4.5 hours in skull bone marrow (SBM) near the superior sagittal sinus (SSS) was found in female and male patients (**a**). Patients with hypertension (**b**) or diabetes mellitus (**c**) exhibit lower percentage changes compared to male and patients without these conditions. Age (**d**) and the Pittsburgh Sleep Quality Index (PSQI) total scores (**e**, n=34) are positively correlated with the percentage change, while the Telephone Montreal Cognitive Assessment (T-MoCA) total scores (**f**, n = 34) was negatively correlated with the percentage change. Each plot displays the sample size, the fitted regression line, and the respective Spearman or Pearson correlation coefficients (ρ or R), with corresponding *P*-values.

**Supplemental Table 1.**
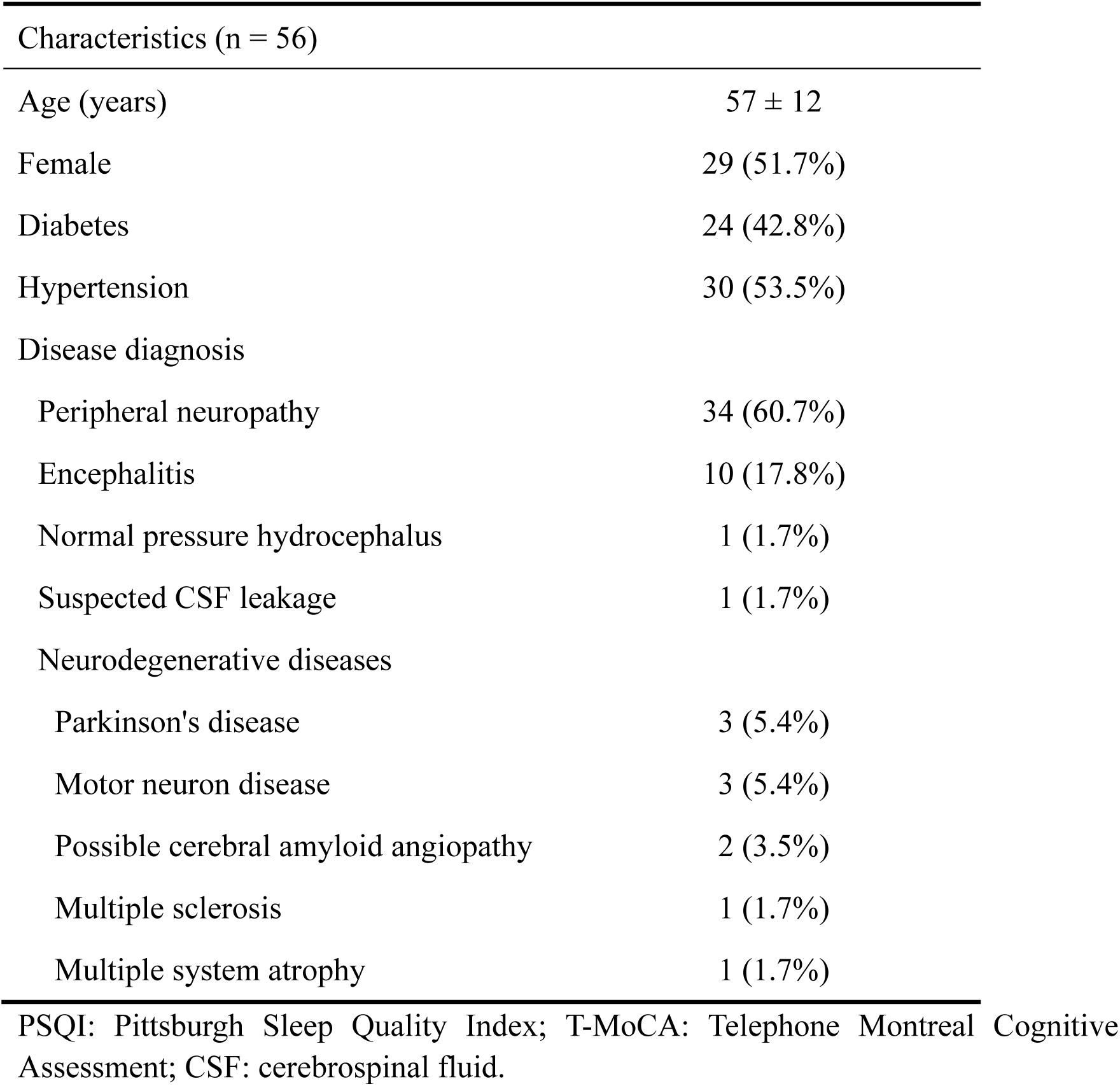
Basic clinical data of patients who completed intrathecal contrast-enhanced MRI, alongside PSQI and T-MoCA tests.

| Characteristics (n = 56) |  |
| --- | --- |
| Age (years) | 57 ± 12 |
| Female | 29 (51.7%) |
| Diabetes | 24 (42.8%) |
| Hypertension | 30 (53.5%) |
| Disease diagnosis |  |
| Peripheral neuropathy | 34 (60.7%) |
| Encephalitis | 10 (17.8%) |
| Normal pressure hydrocephalus | 1 (1.7%) |
| Suspected CSF leakage | 1 (1.7%) |
| Neurodegenerative diseases |  |
| Parkinson's disease | 3 (5.4%) |
| Motor neuron disease | 3 (5.4%) |
| Possible cerebral amyloid angiopathy | 2 (3.5%) |
| Multiple sclerosis | 1 (1.7%) |
| Multiple system atrophy | 1 (1.7%) |

PSQI: Pittsburgh Sleep Quality Index; T-MoCA: Telephone Montreal Cognitive Assessment; CSF: cerebrospinal fluid.

